# An agent-based model of dengue virus transmission shows how multiple uncertainties about vaccine efficacy influence public health impact projections

**DOI:** 10.1101/082396

**Authors:** T. Alex Perkins, Robert C. Reiner, Guido España, Quirine A. ten Bosch, Amit Verma, Kelly A. Liebman, Valerie A. Paz-Soldan, John P. Elder, Amy C. Morrison, Steven T. Stoddard, Uriel Kitron, Gonzalo M. Vazquez-Prokopec, Thomas W. Scott, David L. Smith

**Affiliations:** Department of Biological Sciences and Eck Institute for Global Health, University of Notre Dame, Notre Dame, IN; Fogarty International Center, National Institutes of Health, Bethesda, MD; Department of Epidemiology and Biostatistics, Indiana University, Bloomington, IN; Center for Disease Dynamics, Economics, and Policy, Washington, DC; Department of Entomology and Nematology, University of California, Davis, CA; Department of Global Community Health and Behavioral Sciences, Tulane University School of Public Health and Tropical Medicine, New Orleans, LA; Institute for Behavioral and Community Health, Graduate School of Public Health,San Diego State University, San Diego, CA; Department of Environmental Sciences, Emory University, Atlanta, GA; Institute for Health Metrics and Evaluation, University of Washington, Seattle, WA

## Abstract

Given the limited effectiveness of strategies based solely on vector control to reduce dengue virus (DENV) transmission, it is expected that an effective vaccine could play a pivotal role in reducing the global disease burden of dengue. Of several dengue vaccines under development, Dengvaxia^®^ from Sanofi Pasteur recently became the first to become licensed in select countries and to achieve WHO recommendation for use in certain settings, despite the fact that a number of uncertainties about its profile complicate projections of its public health impact. We used a stochastic, agent-based model for DENV transmission to perform simulations of the public health impact of dengue vaccines in light of two key uncertainties: (1) “statistical uncertainty” about the numerical value of the vaccine’s efficacy against disease, and (2) “biological uncertainty” about the extent to which its efficacy against disease derives from the amelioration of symptoms, blocking of DENV infection, or some combination thereof. Simulations of a generic dengue vaccine showed that the proportion of disease episodes averted following 20 years of routine vaccination of nine-year olds at 80% coverage was sensitive to both the numerical value of vaccine efficacy and to the extent to which efficacy derives from blocking of DENV infection. Simulations of a vaccine resembling Dengvaxia^®^ took into account that vaccine trial results substantially reduced statistical uncertainty but did not address biological uncertainty, resulting in the proportion of disease episodes averted being more sensitive to biological uncertainty than to statistical uncertainty. Taken together, our results indicate limitations associated with the use of symptomatic disease as the primary endpoint of dengue vaccine trials and highlight the importance of considering multiple forms of uncertainty in projections of a vaccine’s public health impact.

## INTRODUCTION

Dengue is the most significant mosquito-borne viral disease of humans, having emerged in recent decades as a major burden on global health [1]. To date, the control and prevention of dengue has relied exclusively on various forms of vector control [2], which have experienced success in certain cases but have not been sufficient to prevent the rapid global expansion of this disease [3]. At the same time, several vaccine candidates for dengue have been under development for a number of years [4], with the Dengvaxia^®^ vaccine from Sanofi Pasteur recently becoming licensed in some countries and recommended for use under specific circumstances by the World Health Organization [5].

A number of concerns about Dengvaxia^®^ arose during clinical trials, the results of which indicated relatively low overall efficacy [6]. In particular, efficacy against disease was significantly lower for children under 9 years of age (44%) compared with children 9 years of age or older (65%). Importantly, the vaccine also appeared to provide higher protection to seropositive than seronegative recipients, especially at young ages (2-5 years). Estimates of vaccine efficacy appeared to further vary among the four dengue virus (DENV) serotypes, with lower efficacy reported for DENV-2 [6]. After accounting for these known effects, we use the term “statistical uncertainty” to refer to residual uncertainty about the numerical value of vaccine efficacy for a given segment of the population exposed to a given DENV serotype.

Another source of uncertainty associated with Dengvaxia^®^ has to do with the clinical nature of the trial endpoints. Specifically, the primary endpoint for efficacy trials of Dengvaxia^®^ was virologically confirmed dengue among trial participants who experienced acute febrile illness; i.e., a fever of ≥38 °C for at least two consecutive days [6]. This choice of endpoint is potentially problematic, because a large proportion of DENV infections result in either mild symptoms or no detectable symptoms whatsoever [7] in people that are nonetheless capable of infecting mosquitoes [8]. Consequently, it is unclear whether Dengvaxia^®^ confers protection to individuals who do not experience an acute febrile illness. Among trial participants for whom acute febrile illness was averted due to vaccination, the vaccine could have either (1) blocked DENV infection altogether, or (2) ameliorated symptoms but still allowed for infection and onward transmission. We use the term “biological uncertainty” to refer to this combination of unknown factors.

In addition to fundamental challenges that biological uncertainty poses for estimating vaccine efficacy from trial data [9], it has substantial implications for projections of the public health impact of vaccination. In general, the public health impact of any vaccine derives from three sources. First, vaccine recipients experience direct protection in the form of reduced incidence of disease. Second, unvaccinated individuals experience indirect protection through the buildup of herd immunity in vaccine recipients, which prevents them from becoming infected and contributing to onward transmission in the future. Third, unvaccinated individuals experience indirect protection if vaccination blocks infection in a vaccinated individual “upstream” of them in a transmission chain. Projections of public health impact that capture the first two of these effects can be made based on vaccine efficacy estimates from clinical trials and knowledge of vaccination coverage. Projections that capture the third of these effects depend additionally on understanding the extent to which vaccination blocks infection.

To assess the influence of these uncertainties on public health impact projections, we simulated DENV transmission in the presence and absence of routine vaccination under a range of assumptions about statistical and biological uncertainty. In addition to simulations using a vaccine parameterized with Dengvaxia^®^ trial results, we performed simulations using a hypothetical and more generic dengue vaccine to explore the implications of different forms of uncertainty for forthcoming dengue vaccines. We performed these simulations with an agent-based simulation model of DENV transmission developed and calibrated for the city of Iquitos, Peru, which has had ongoing studies of dengue epidemiology for nearly two decades [10,11]. One advantage of our model relative to existing simulation models of DENV transmission [12,13] is that it features the most realistic model available for fine-scale human movement in a dengue-endemic area [14], which is important for realistic quantification of the extent to which blocking of DENV infections in vaccine recipients may disrupt transmission. The model also features human demographic rates consistent with U.N. estimates [15] and accounts for interannual variability in serotype dynamics in a manner consistent with longitudinal studies in Iquitos over an 11-year period [16], making it suitable for generating long-term projections of vaccine impact.

## METHODS

### Model overview

We developed a stochastic, agent-based model for simulating DENV transmission that is parameterized in a number of respects around studies of dengue epidemiology conducted in Iquitos, Peru. The model simulates DENV transmission in a population of approximately 200,000 people residing in the core of Iquitos, which consists of 38,835 geo-referenced houses and 2,004 other buildings [17]. Events such as mosquito biting, mosquito death, and movements by humans and mosquitoes are scheduled to occur at continuous time points throughout the day (Fig. 1), with updating of individuals’ statuses with respect to infection, immunity, and demographics occurring once daily. We described the model in full detail in Supporting Information 1, following the ODD (Overview, Design concepts, Details) Protocol [18,19] for describing agent-based models. In the three paragraphs below, we provide an overview of key features of the model pertaining to humans, mosquitoes, and viruses, respectively.

**Figure 1.**
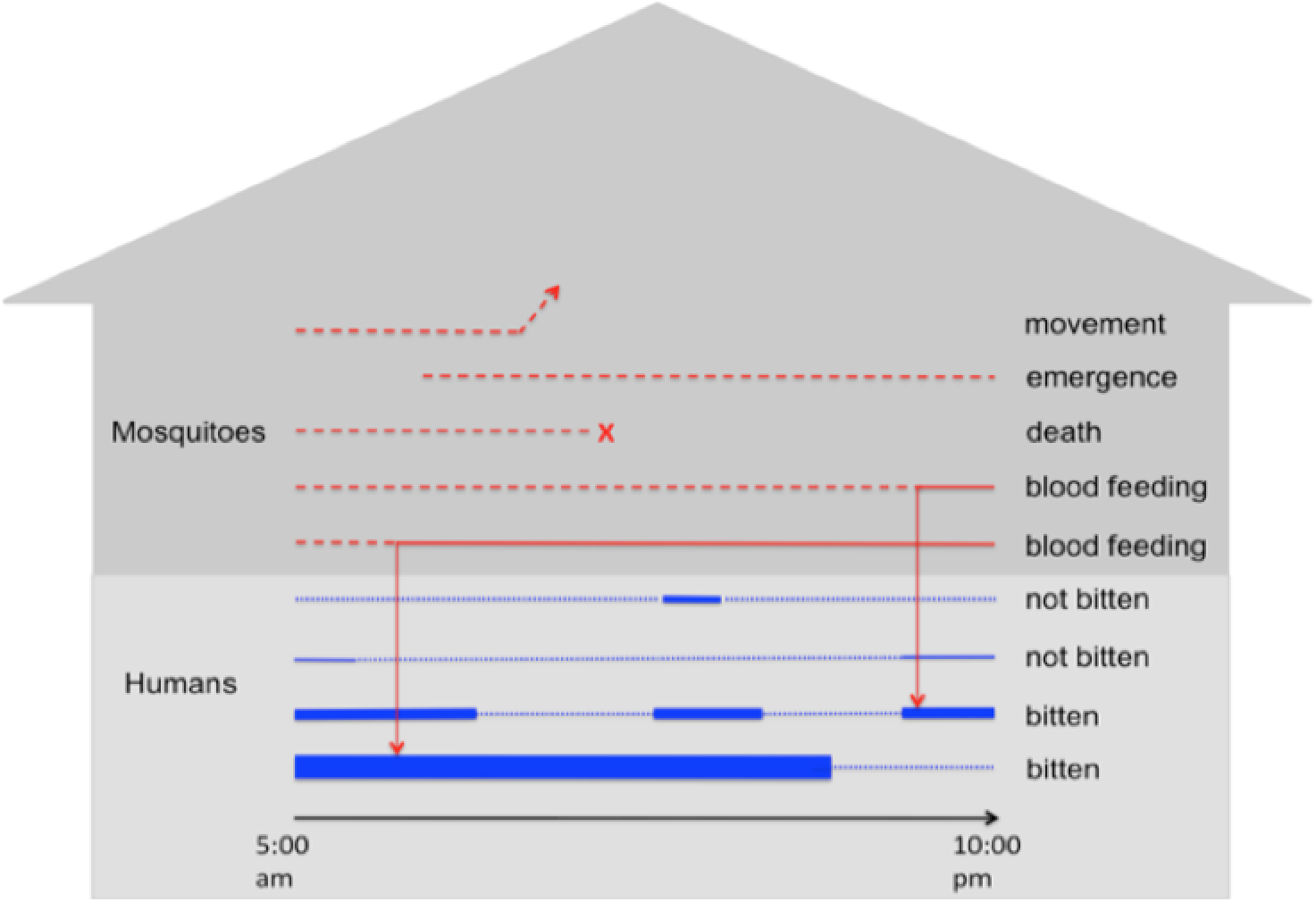
Example of events that occur over the course of a single day at a single location. Red lines correspond to individual mosquitoes, with dashed and solid lines representing host-seeking and resting states, respectively. Blue lines refer to individual people, with thin dotted lines indicating that the person is at another location at that time and thick solid lines indicating their presence at the location at that time. The thickness of the solid blue lines indicates the relative attractiveness of each person to blood feeding by mosquitoes.

Humans are populated in the city consistent with national age and sex distributions for Peru and in individual houses consistent with demographic data collected over the course of studies in Iquitos. Birth and death processes are parameterized consistent with demographic estimates and future projections by the United Nations [15], with age- and year-specific death rates, year-specific population birth rates, and age- and year-specific relative fertility among females aged 15-49. Aging involves the acquisition of lifelong, serotype-specific immunity as each person is exposed, and sex-specific growth of an individual’s body size over the course of childhood to allow for an effect of body size on propensity to be bitten by mosquitoes [20]. Each individual human possesses a unique “activity space,” which is defined as an average pattern of time allocation across all the locations that they frequent between 05:00 and 22:00, when risk of biting from *Ae. aegypti* mosquitoes is expected to be highest [21]. Individuals move about this activity space in a manner based on retrospective interviews performed on residents of Iquitos and modeled in a way described previously by Perkins et al. [14].

The number of adult female mosquitoes in the area is determined by a combination of mosquito emergence and death processes. Mosquito death occurs according to unique daily, temperature-dependent rates derived from Brady et al. [22]. Mosquito emergence occurs differentially by location according to unique daily emergence rates that were estimated by determining what emergence patterns, when combined with the death process in our model, would yield spatio-temporal patterns of mosquito density consistent with estimates by Reiner et al. [23]. In addition to emergence and death, mosquitoes move from their current location to a nearby location on any given day with a fixed probability [24]. They engage in biting at temperature-dependent rates that differ depending on whether it is the mosquito’s first bite [25] or a subsequent bite [26]. They select an individual on whom to blood feed based on the body size of each person present at a location at the time that a mosquito bites [14,20]. Because the emphasis of the present analysis is on vaccination rather than vector control, we deferred the inclusion of additional entomological details for future work.

The model allows for the transmission of all four DENV serotypes, which are assumed to be identical in the following respects based on empirical studies: infectiousness [8,27], extrinsic and intrinsic incubation periods [28], and rate of symptomatic disease [29]. For infectiousness of mosquitoes to people, we adopted a value used by another modeling study (0.9, [30]), given the difficulty of estimating this parameter empirically. For infectiousness of people to mosquitoes, we employed a time-varying function of infectiousness based on empirical data [31]. Individual people can experience up to four distinct infections over the course of their lifetimes, as they experience lifelong immunity to each serotype to which they have been exposed and temporary cross-immunity to all serotypes following exposure [32]. Viruses of each serotype are seeded into the population at a time-varying rate through infected people that are each simulated to have an activity space identical to a randomly chosen resident for the duration of their infection. Because currently available data in the geographic information system only permits us to simulate approximately half the population of the metropolitan area, we viewed these infections in temporary individuals primarily as representative of people from other parts of the city moving DENV into the population represented explicitly in our model. Were we to model the entire population of Iquitos, we expect that fewer such infections would be necessary to seed transmission within our synthetic population. In addition, although Iquitos is relatively isolated in general, some limited importation of DENV is known to occur from surrounding areas [33].

### Model calibration

Wherever possible, we parameterized the model directly based on empirical estimates from Iquitos or from studies conducted elsewhere and reported in the literature. This included human demography [15,20], human mobility [14], human-mosquito encounters [20], mosquito abundance patterns in space and time [34–36], mosquito movement [24], mosquito mortality [22], mosquito blood-feeding rates [25,26], virus incubation in mosquitoes and humans [28], infectiousness of humans to mosquitoes [31], and naturally acquired immunity to DENV [32].

Two primary uncertainties that we were not able to quantify a priori were DENV importation into Iquitos and the scaling relationship between household mosquito surveys and true mosquito abundance. We calibrated those parameters such that simulated model behavior was consistent with empirical estimates of time-varying, serotype-specific patterns of incidence of DENV infection [16]. We did so on the basis of the goodness of fit of simulated incidence *I*_*s,t*_ of serotype *s* at time *t* to probabilistic estimates of *I*_*s,t*_ by Reiner et al. [16]. For each month between January, 2000 and June, 2010, we first performed maximum-likelihood fitting of a Dirichlet distribution to 1,000 draws of the serotype proportions of *I*_*s,t*_ and a normal distribution to 1,000 draws of the total incidence *I*_*t*_ at time *t* from the posterior distribution estimated by Reiner et al [16]. We then used the product of the probability densities of those distributions evaluated at the simulated values of *I*_*s,t*_ for each serotype as our measure of goodness of fit.

Using this measure of goodness of fit, we obtained estimates of unknown parameters for DENV importation and scaling of mosquito abundance using a particle filtering algorithm. The premise of this algorithm is to make use of the fact that most of the unknown parameters pertain to only a portion of the time series -- and thereby only a portion of the likelihood -- to allow for calibrating different subsets of the unknown parameters sequentially rather than simultaneously. There are a wide range of particle filtering algorithms, but ours most closely resembles a sequential importance resampling algorithm [37].

The first step in our algorithm involved proposing a set of 1,000 initial particles spanning a range of parameter values, simulating the first year of the model for each particle, evaluating the goodness of fit measure described above on a monthly basis within the first year, and combining the monthly goodness of fit measures to obtain an annual goodness of fit measure for each particle for the first year. Next, we resampled the particles 1,000 times with replacement weighted by

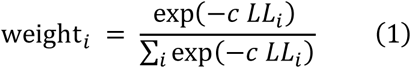

for each particle *i*, where *c* is a scaling parameter that we tuned to a value of 0.1 to result in resampled particles containing at least 10% of the original particles. We then obtained maximum-likelihood estimates of the means and covariance matrix describing the distribution of the particles. Using that fitted multivariate normal distribution, we then drew 1,000 new particles and simulated both the first and second year of transmission. We then computed the goodness of fit measure for the first two years by aggregating monthly goodness of fit measures across the first two years. Additional steps in the algorithm were repeated in the same way in yearly increments through the last year for which empirical estimates of time-varying, serotype-specific incidence were available. Finally, we performed two additional rounds of resampling on the full time series following the last year of simulation and particle resampling. The resulting set of 1,000 particles constituted our distributional estimate of parameter values most consistent with available empirical estimates.

### Vaccine efficacy

#### Dengvaxia^®^-like vaccine

The mode of action of the Dengvaxia^®^ vaccine is not completely clear, given that there are multiple mechanisms by which results from clinical trials could have come about [38]. Although there is evidence that vaccine efficacy differs by serotype [6], no data have been published to date that indicate how serotype-specific effects might manifest differently in individuals of different ages and serostatuses (i.e., seropositive for seronegative for past DENV exposure). Because serostatus-specific effects are most concerning from a safety perspective [39] and because of clear interactive effects between age and serostatus [6], we prioritized those effects and assumed in the model that efficacy applies equally to each of the four serotypes.

We modeled vaccine efficacy against disease (i.e., the primary trial endpoint) as a function of age and serostatus, which is consistent with one hypothesis for how the vaccine achieves its efficacy [6,38]. Specifically, for a given serostatus, we modeled the relationship between age and vaccine efficacy (VE) against disease as

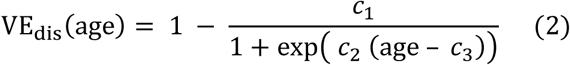

using serostatus-specific values of *c*_1_, *c*_2_, and *c*_3_. This functional form was chosen based on the fact that its shape is relatively flexible and due to the fact that it yields a monotonic relationship between age and VE_dis_. To obtain point estimates of *c*_1_, *c*_2_, and *c*_3_ for both seropositive and seronegative vaccine recipients, we fitted eqn. (2) under different values of these parameters to mean estimates of VE_dis_ for 2-9 and 10-16 year olds reported in Fig. 2 of Hadinegoro et al. [6] on the basis of least squares using the Nelder-Mead algorithm as implemented in the optim function in R [40]. These calculations assumed an even age distribution within each age class in the trials.

**Figure 2.**
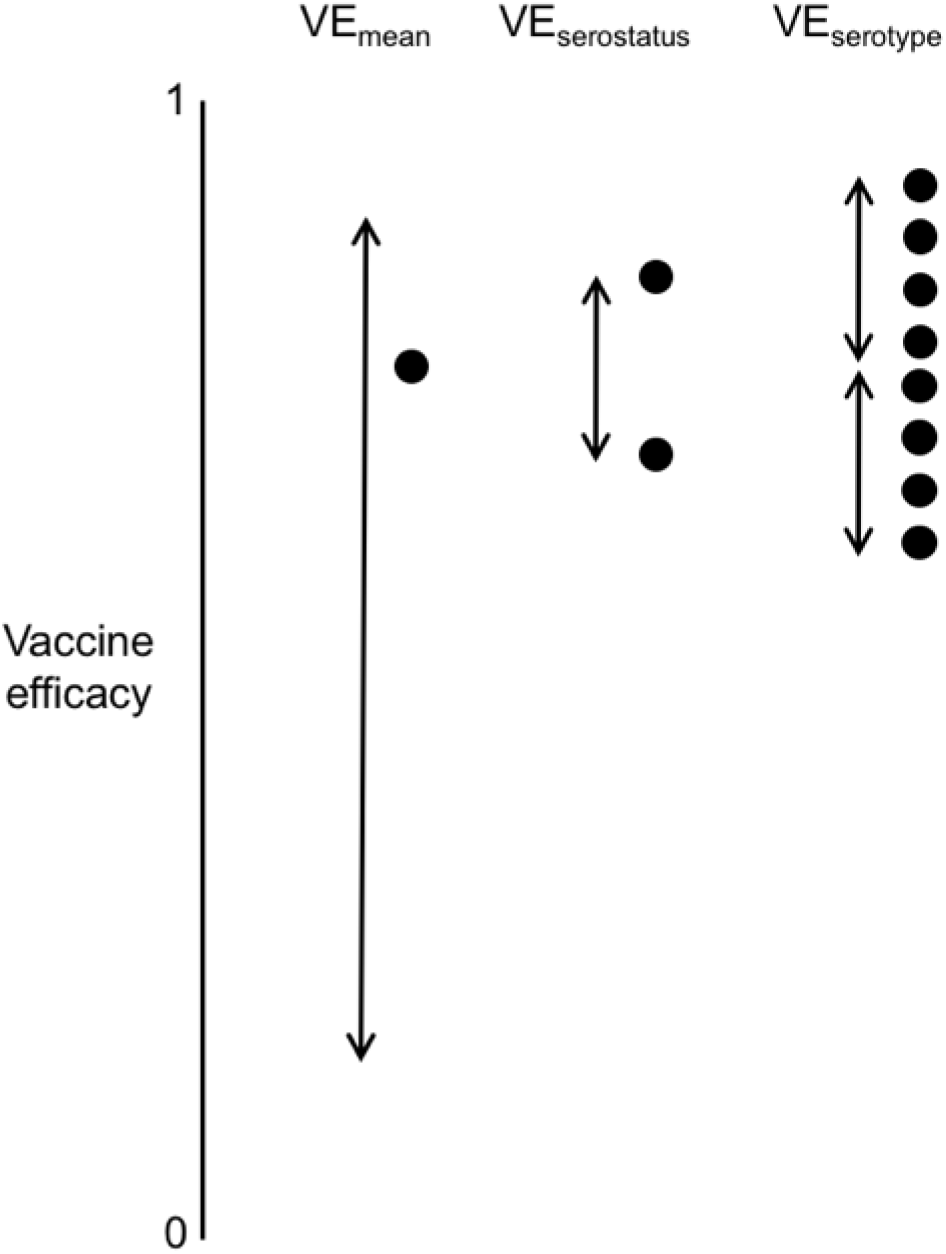
Parameters used to define the profile of a hypothetical, generic dengue vaccine. The eight dots on the right correspond to eight distinct values of vaccine efficacy (VE) that apply to people with 8 possible combinations of serostatus (seropositive or seronegative) and exposure at the time of vaccine protection to one of the four DENV serotypes. The mean VE across all eight combinations is defined by VE_mean_, with VE_serostatus_ determining the increment up (for seropositives) or down (for seronegatives) for serostatus and 2 × VE_serotype_ determining the range of serotype-specific VE for a person with a given serostatus. The ranking of serotype-specific VE among the four serotypes is randomized across simulations.

We modeled statistical uncertainty around estimates of VE_dis_ with a parameter σthat describes the standard deviation of the log of the relative risk (RR), defined mathematically as

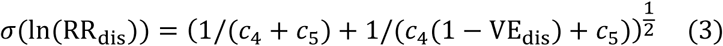

to yield a one-to-one relationship between VE_dis_ and the standard deviation of the log of RR_dis_. To fit values of *c*_4_ and *c*_5_, we used a method based on the assumption of asymptotic normality of the log of the ratio of Poisson rates [41], applied to standard errors presented in Fig. 2 of Hadinegoro et al. [6]. To then take a random draw of VE_dis_ for a given instance of the simulation with statistical uncertainty consistent with that reported by Hadinegoro et al. [6], we drew a normal random variable with mean 0 and standard deviation 1, multiplied it by σadded the result to ln(1-VE_dis_), exponentiated the result, and subtracted it from 1 [6].

In all simulations, we applied the same standard normal random draw to the calculation of VE_dis_ for both seropositive and seronegative vaccine recipients. We assumed that the vaccine is leaky, meaning that protection from infection is not perfect; i.e., an individual has some chance of becoming infected each time they are exposed and has some chance of developing disease each time they are infected. The chances of those events occurring in a vaccine recipient are lowered proportional to relative risk: RR_inf|exp_ and RR_dis|inf_, respectively (Table 1).

**Table 1.** Definitions of key terms.

| Term | Symbol | Definition |
| --- | --- | --- |
| Relative risk of disease conditional on infection | $RR_{dis\vert inf}$ | Proportion of vaccine recipients that experience disease after becoming infected relative to the proportion of placebo recipients that experience disease after becoming infected. |
| Relative risk of infection conditional on exposure | $RR_{inf\vert exp}$ | Proportion of vaccine recipients that become infected after being bitten by an infectious mosquito relative to the proportion of placebo recipients that become infected after being bitten by an infectious mosquito. |
| Relative risk of disease | $RR_{dis}$ | Proportion of vaccine recipients that experience disease after being bitten by an infectious mosquito relative to the proportion of placebo recipients that experience disease after being bitten by an infectious mosquito. This is equal to the product of $RR_{dis\vert inf}$ and $RR_{inf\vert exp}$. |
| Vaccine efficacy against disease | $VE_{dis}$ | $1 - RR_{dis}$ |
| Proportion of protection against disease derived from protection against infection | $p$ | This parameter relates $RR_{inf\vert exp}$ to $RR_{dis}$ according to the relationship $RR_{inf\vert exp} = RR_{dis}^p$. Likewise, it is implied that $RR_{dis\vert inf} = RR_{dis}^{1-p}$ and $RR_{dis} = RR_{dis\vert inf} \times RR_{inf\vert exp}$. |
| Quantile of $RR_{dis}$ estimate | $q$ | Quantile between 0 and 1 applied to the uncertainty distribution of the $RR_{dis}$ estimate. |

#### Generic dengue vaccine

To examine the robustness of our results to assumptions about the vaccine profile of Dengvaxia^®^, we also considered a hypothetical and more generic dengue vaccine. We characterized this vaccine’s profile with three parameters: VE_mean_, VE_serostatus_, and VE_serotype_ (Fig. 2). Under this model, an individual’s RR depends on their pre-vaccination serostatus and the DENV serotype to which they were exposed, but not their age. To ensure that VE_mean_ does in fact represent a mean, each individual’s VE begins there and is adjusted up or down by VE_serostatus_ and VE_serotype_. VE_serostatus_ is always subtracted from VE_mean_ for seronegative individuals and added for seropositives. From there, an increment is added or subtracted such that the four serotype-specific VEs span a range of 2 × VE_serotype_. Which serotypes have higher or lower VE is randomized across simulations. The RR of an individual is calculated as 1 less the VE determined by their serostatus and serotype, and RR_dis|inf_ and RR_inf|exp_ are calculated the same as for the Dengvaxia^®^-like vaccine.

### Vaccine impact projections

For the Dengvaxia^®^-like vaccine, we performed 1,000 pairs of simulations with parameter particles drawn from the posterior and *p* (representing “biological uncertainty”) and *q* (representing “statistical uncertainty”) sampled between values of 0 and 1 with the sobol function in the pomp package [42] in R [40] to maximize coverage of the *p*-*q* parameter space. For the generic vaccine, we performed 1,000 pairs of simulations with parameter particles for *p*, VE_mean_, VE_serostatus_, and VE_serotype_, sampling across respective ranges of 0-1, 0.15-0.85, 0-0.15, and 0-0.15, also using the sobol function. Values of the latter three parameters were chosen to ensure that the maximum VE could not exceed 1 or fall below 0 and that a broad range of VE_mean_ was covered.

The two simulations in each pair exhibited identical dynamics for the first 11 years, because they were both driven by the same parameter particle and both shared common random number seeds for processes related to mosquito-human contact and DENV infection, respectively. Following that initial time period, we continued one simulation without vaccination but commenced the other with routine vaccination at age nine, both for an additional 20 years. Consistent with other model-based projections of Dengvaxia^®^ public health impact [43], we assumed 80% coverage. For each simulation pair, we recorded the following in the population as a whole: proportion of cumulative infections averted and proportion of cumulative disease episodes averted, with both accruing over the period that followed the time period calibrated to Iquitos.

## RESULTS

### Model calibration

The behavior of the calibrated model was largely in agreement with the estimates of time-varying, serotype-specific incidence of infection to which it was calibrated (Fig. 3). For all serotypes, the 95% prediction interval of simulations from our agent-based model and estimates from Reiner et al. [16] overlapped for the majority of the 2000-2010 timeframe. Both patterns reflected relatively low and seasonally variable patterns of DENV-1 and DENV-2 transmission, and both captured two relatively large seasonal peaks in 2002-2003 for DENV-3 and in 2009-2010 for DENV-4, coinciding with the respective invasions of those serotypes. These results were nearly identical when the model was calibrated under different assumptions about the duration of temporary cross-immunity (Figs. S1-S4), indicating that the model’s ability to reproduce dynamics from Iquitos is robust to this assumption and that our algorithm for calibrating the model led to convergent estimates across multiple runs.

**Figure 3.**
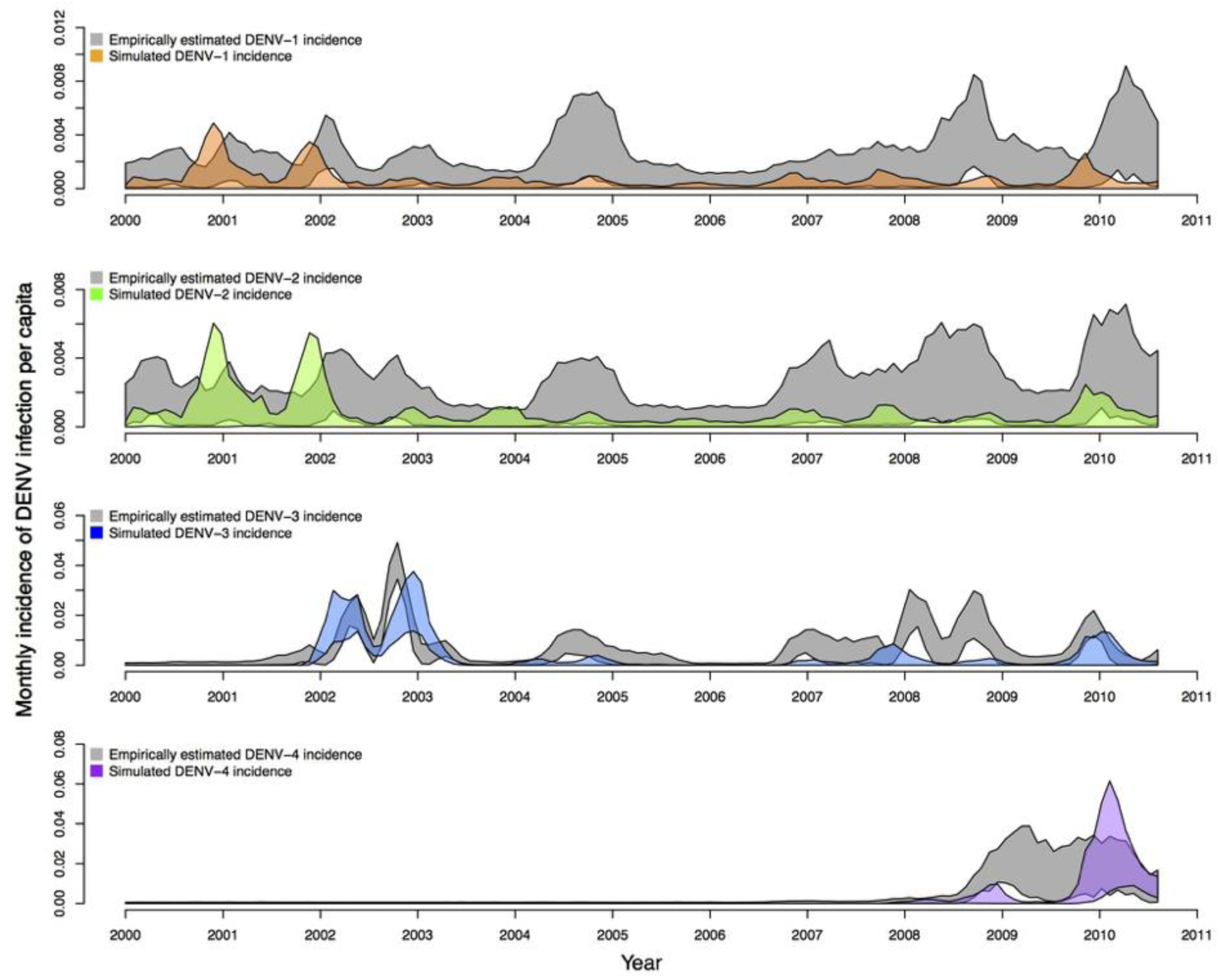
Monthly, serotype-specific incidence of infection per capita, as estimated by Reiner et al. [16] (gray bands) and as reproduced by our calibrated model (colored bands). Bands show the range of values in which 95% of simulated values lie for a given serotype in a given month. These values were obtained under the assumption that the period of temporary cross-immunity is exponentially distributed with a mean of 686 days.

Although the calibrated model was in relatively good agreement with estimates of incidence patterns by Reiner et al. [16] for large transmission seasons early in the occurrence of a given serotype, the model had a tendency to produce somewhat lower incidence patterns afterwards (Fig. 3). These periods of lower incidence were associated with lower population susceptibility to a given serotype (Fig. 4) and tended to require a larger number of infections to seed transmission (Fig. 5). The relatively low number of infections required to seed the larger epidemics was encouraging with regard to the model’s ability to reproduce large epidemics in Iquitos, but the relatively high number of infections required to seed inter-epidemic transmission suggests that the model may not be ideal for testing hypotheses about the factors that enable DENV to persist in this population over extended periods of time.

**Figure 4.**
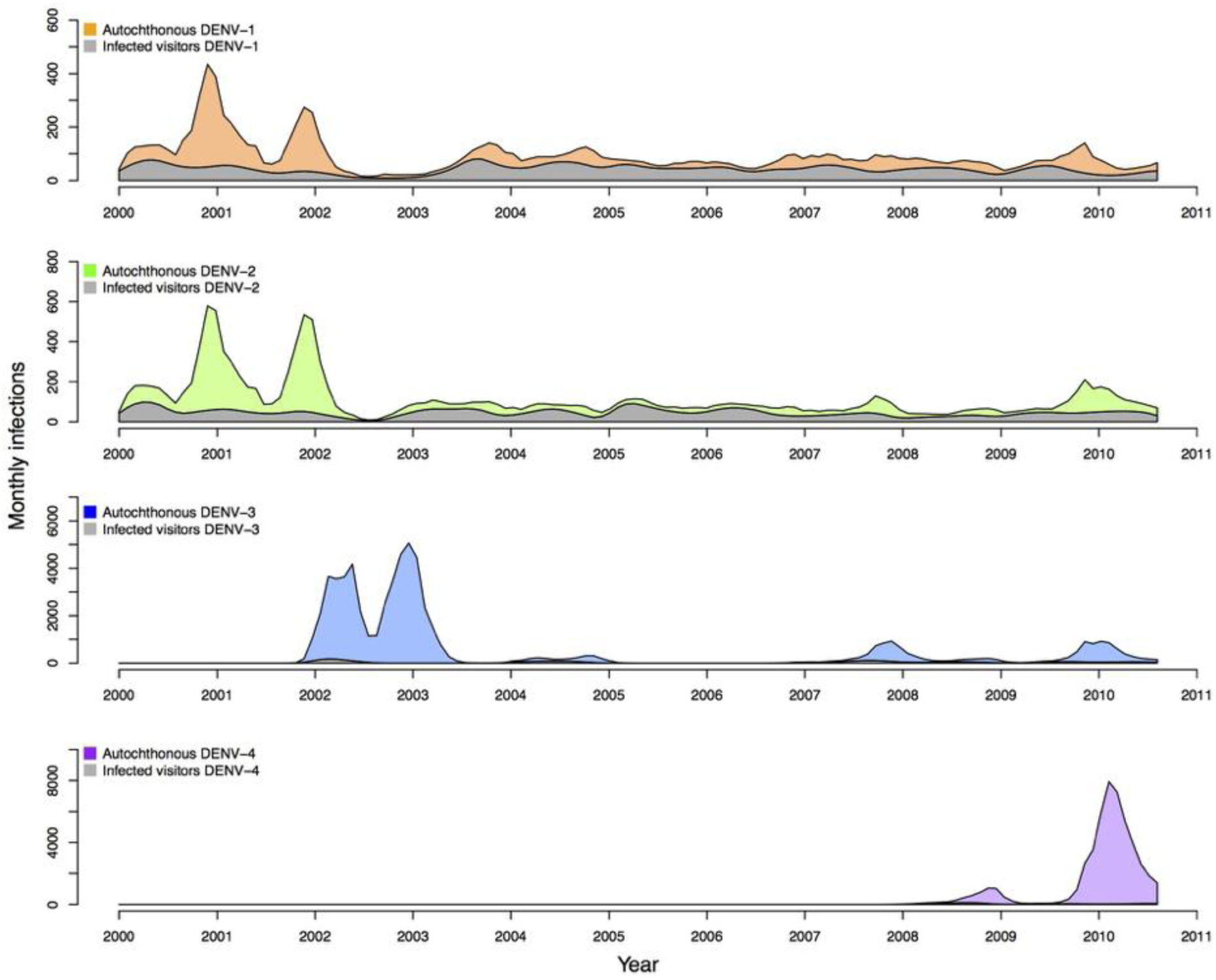
Median numbers of infections on a monthly basis for each serotype, separated by whether the infection was acquired through biting by an infectious mosquito (colored) or by exogenously driven infections (gray) that were used to seed transmission in the model. These bands represent median values across the set of calibrated parameter values, and the colored bands are added on top of the gray bands. These values were obtained under the assumption that the period of temporary cross-immunity is exponentially distributed with a mean of 686 days.

**Figure 5.**
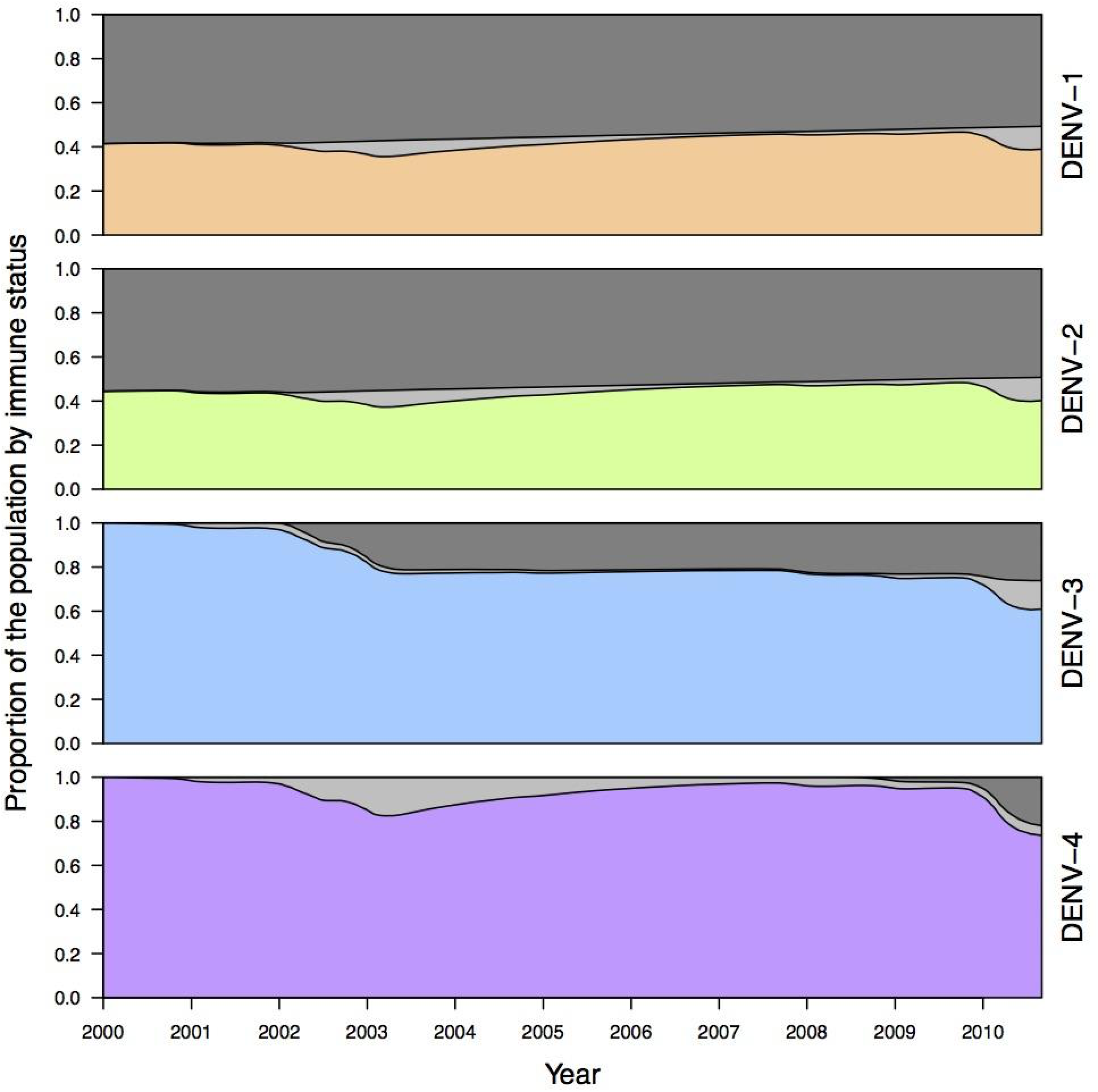
Proportion of the population during the period of the calibration that was not immune to a given serotype (colored), temporarily cross-immune to that serotype (light gray), or permanently immune to that serotype (dark gray). Values shown reflect medians across 1,000 simulations drawing parameters from across the set of particles obtained through the calibration process, under the assumption that the period of temporary cross-immunity is exponentially distributed with a mean of 686 days. Variability around these values was similar in scale to that shown in Fig. 3.

### Vaccine efficacy

We obtained estimates of the parameters for relative risk in eqn. (2) that best match empirical estimates [6] of *c*_1_ = 0.47, *c*_2_ = 0.148, and *c*_3_ = 9.17 for seropositive vaccine recipients and *c*_1_ = 1.26, *c*_2_ = 0.28, and *c*_3_ = 9.27 for seronegative vaccine recipients. We obtained estimates of the parameters determining the standard error of the log of the risk ratio in eqn. (3) of *c*_4_ = 100 and *c*_5_ = 0.5. Under this model and with these parameters, relative risk decreased steeply with age until around age 20, when it began to decrease more slowly towards almost no risk in older people (Fig. 6). As in the clinical trial data, relative risk under our model was several fold lower in seropositive than seronegative children, and relative risk in excess of 1 was likely only at ages well below nine years (Fig. 6).

**Figure 6.**
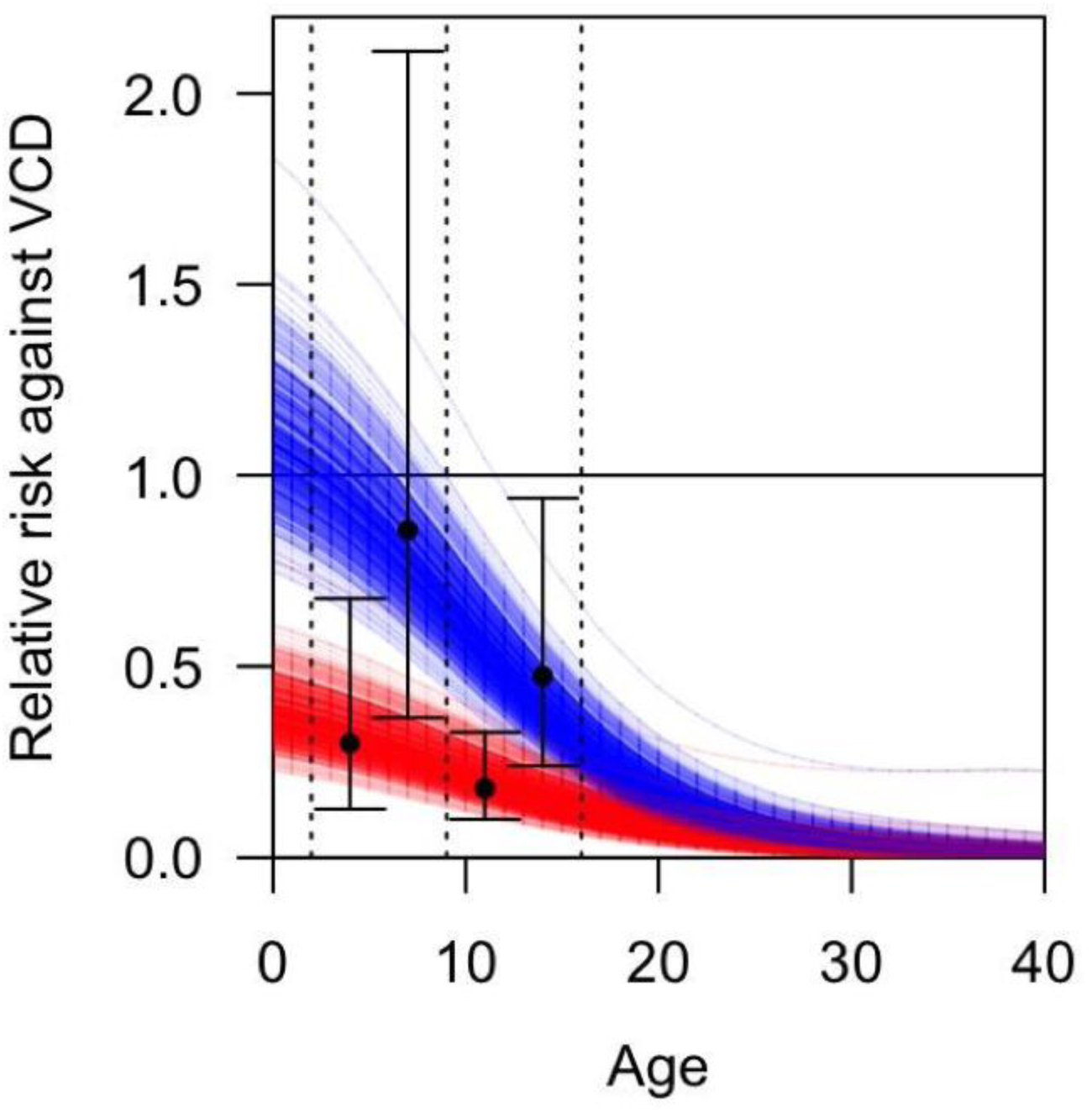
Relative risk of disease, RR_dis_, as a function of age and serostatus (blue = seronegative, red = seropositive) estimated from vaccine trial data for the endpoint of virus-confirmed disease (VCD) [6]. Each line represents a distinct random draw from the distribution of these relationships. Black dots correspond to point estimates of relative risk of disease in the trial for a given age group (2-9 left, 9-16 right) and serostatus (red vs. blue), and error bars indicate 95% confidence intervals around those estimates.

Implementing either vaccine scenario in our simulations required an estimate of RR per event, rather than RR over the course of a trial [6]. For a leaky vaccine, these two different interpretations of RR may vary depending on how many times study participants are exposed [44]. In Supporting Information 2, we showed that these values of RR are unlikely to differ for a dengue vaccine by more than 5%. Given that relatively small difference and in the absence of more detailed information about the number of exposures that participants experienced during Dengvaxia^®^ trials, we operated under the assumption that these two interpretations of RR were equal.

Under our assumptions about how efficacy observed in trials derived from two separate types of protection, an assumption of equal parts protection against infection and protection against disease (i.e., *p* = 0.5) gave, on average, relative risks of 48% for either infection or disease in seropositive nine-year olds and 80% for either in seronegative nine-year olds (Fig. 7). In the event that 90% of protection derived from protection against disease and only 10% from protection against infection (i.e., *p* = 0.1), the relative risk for seropositive nine-year olds was 27% for disease and 87% for infection and 68% for disease and 96% for infection for seronegative nine-year olds (Fig. 7).

**Figure 7.**
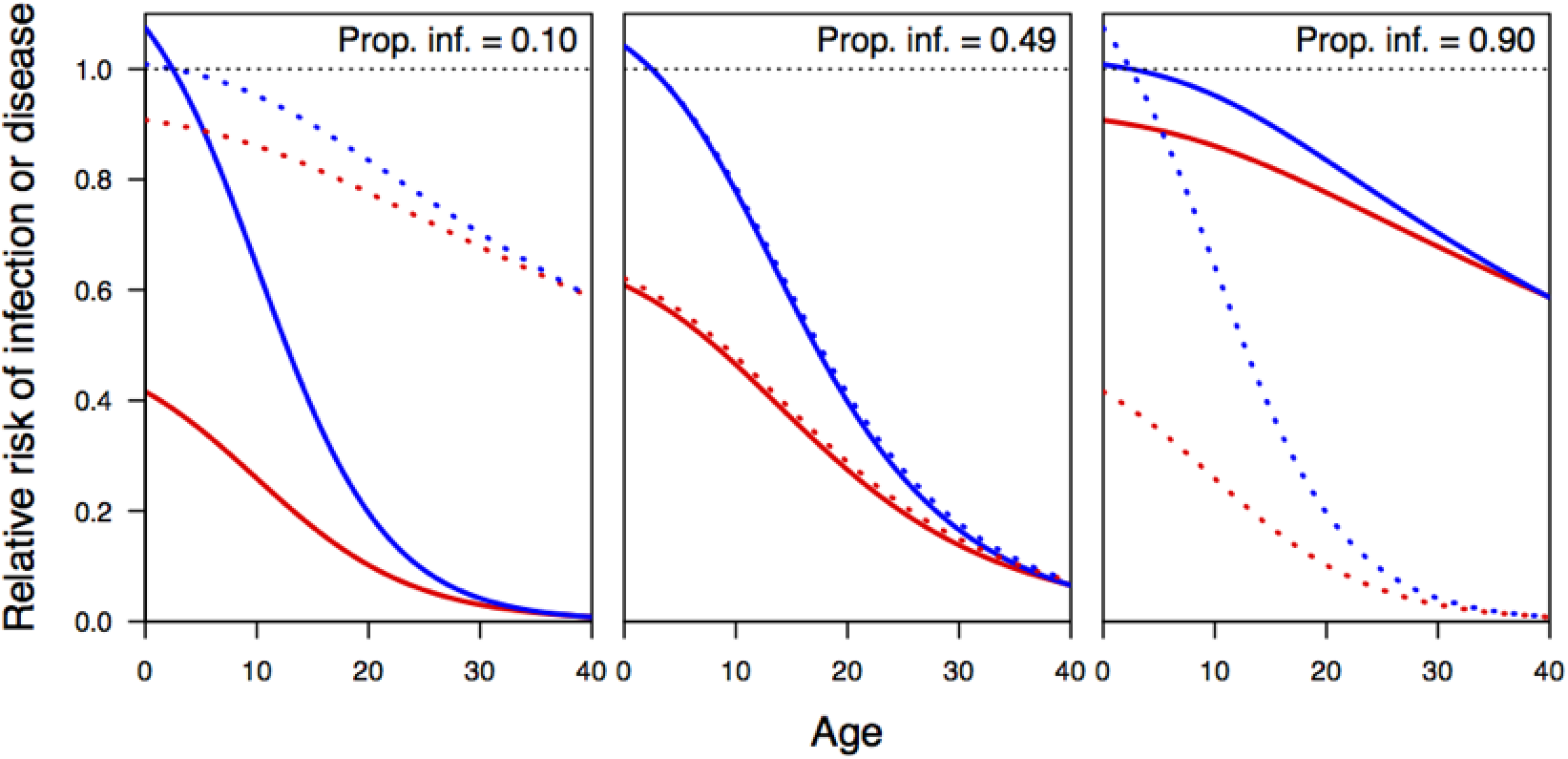
Relative risk of infection conditional on exposure (dashed) and of disease conditional on infection (solid) for seropositive (red) and seronegative (blue) individuals of different ages. These relationships are shown for three different values of the parameter *p* that specifies the proportion of the overall efficacy against disease that is attributable to protection against infection conditional on exposure.

### Vaccine impact projections

Following the calibration period, pairs of simulations were run for 20 additional years, with one simulation in each pair implementing routine vaccination at age nine and one not. The extent of differences between paired simulations depended on the extent to which vaccine efficacy derived from protection against infection or protection against disease. In three pairs of simulations in which the Dengvaxia^®^-like vaccine derived its efficacy completely from protection against infection (*p*=1), epidemic size was noticeably smaller during outbreak years in simulations with vaccination (Fig. 8, green lower than black in left column). In three pairs of simulations in which the vaccine derived its efficacy completely from protection against disease but not infection (*p*=0), epidemic size was somewhat lower in terms of incidence of disease but essentially identical in terms of incidence of infection (Fig. 8, green similar to black in right column). Due to extensive nonlinearities associated with DENV transmission dynamics, these effects were not uniform in time but instead had complex effects on epidemic size, despite controlling for differences associated with random number seeds to the greatest extent possible. The quantitative extent of these differences for these few examples should not be emphasized; instead, trends across the simulation sweeps as a whole are more revealing.

**Figure 8.**
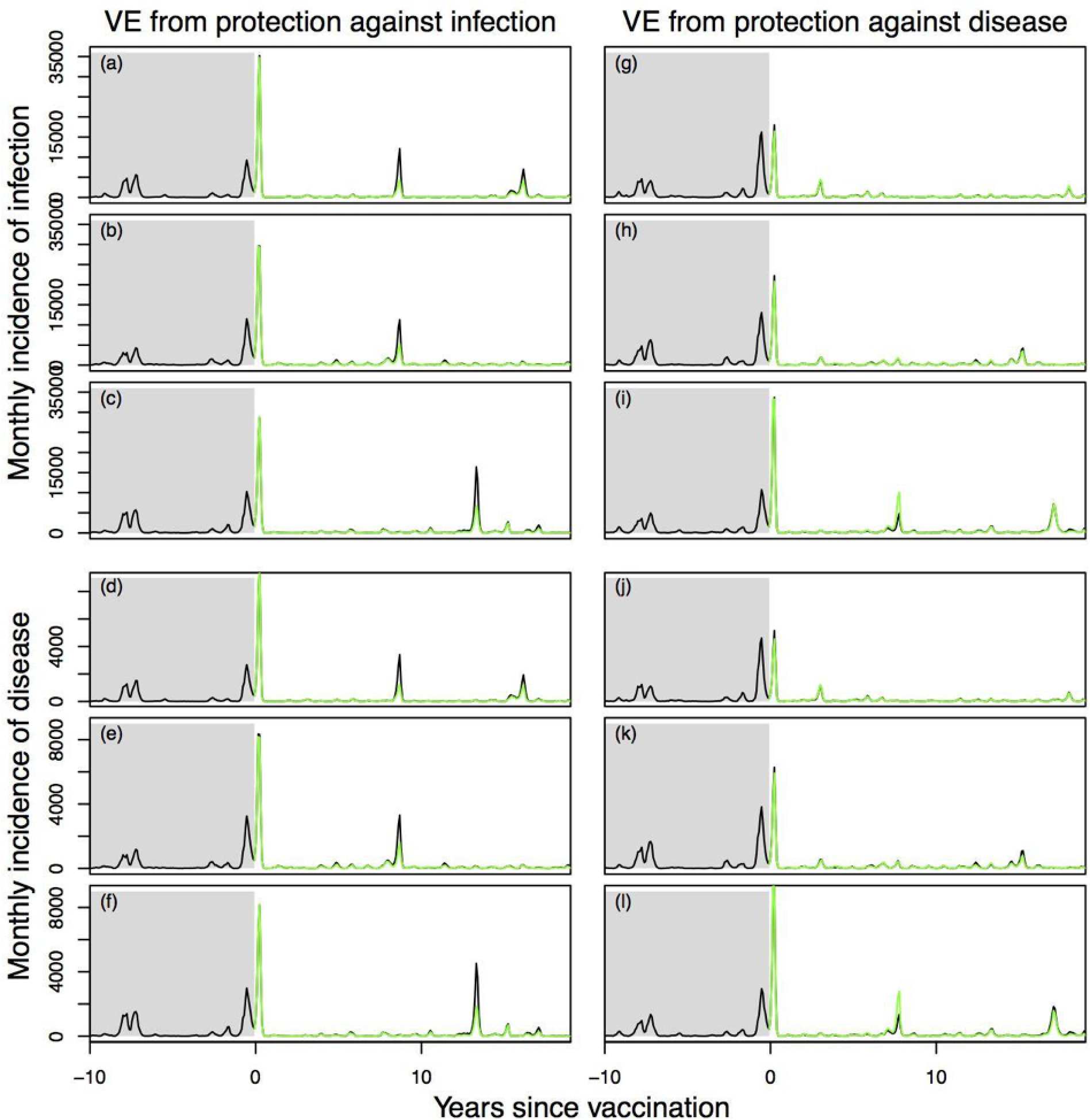
Example time series of annual incidence of human DENV infections simulated from the model with vaccination (green) and without (black). Each panel shows a pair of simulations with two common random number streams for (i) events related to the demography, movement, and mixing of mosquitoes and people, and (ii) events related to infection and disease. Prior to year 0, both simulations in each pair are identical and follow dynamics calibrated as shown in Fig. 3. Beginning in year 0, routine vaccination commences in the simulation colored in green, but not in the one in black. Three different realizations are shown for each of two scenarios (left: *p* = 1; right: *p* = 0).

Simulations conducted with the Dengvaxia^®^-like vaccine across the full range of the parameter *p* showed that, on average, the proportion of cumulative infections averted and the proportion of cumulative disease episodes averted were both maximized (at 14.3% and 13.1%, respectively) when vaccine efficacy derived fully from protection against infection (*p*=1) (Fig. 9, left). Under the opposite extreme (*p*=0), a lower proportion of cumulative disease episodes were averted (4.9%), reflecting direct protection of individuals who received the vaccine. The proportion of cumulative infections averted was approximately zero under this scenario, as infections experienced neither by vaccine recipients nor by others were prevented by a vaccine that had no impact on the ability of vaccine recipients to transmit DENV upon becoming infected. Differences in the proportion of cumulative infections averted and the proportion of cumulative disease episodes averted were not as large across the range of *q* (Fig. 9, right). Across the full range of vaccine efficacy quantiles captured by *q*, the proportion of cumulative infections averted varied by 0.87% (Fig. 9, right). Results for cumulative disease episodes averted were similar (0.81%).

**Figure 9.**
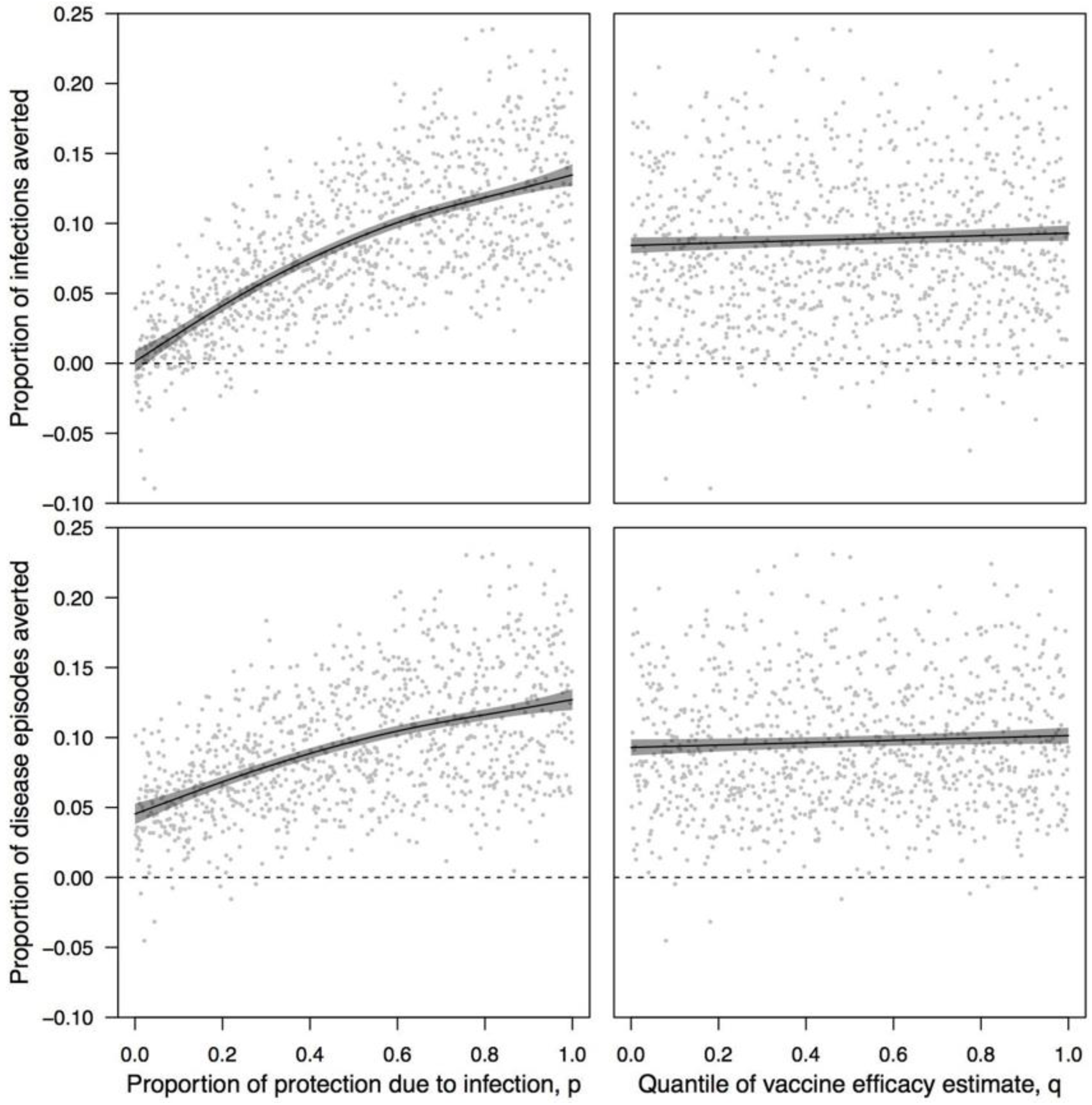
Impacts of vaccination assessed in 1,000 pairs of simulations with and without vaccination with the Dengvaxia^®^-like vaccine, with simulation pairs varying with respect to the proportion of vaccine efficacy due to protection from infection, *p*, (left column) and the quantile of estimated vaccine efficacy, *q* (right column). The proportion of cumulative infections (top row) and disease episodes (bottom row) averted were calculated as the number of each in the simulation without vaccination minus the number of each in the simulation with vaccination, both following 20 years of routine vaccination of 9-year olds at 80% coverage. Lines show the proportion of infections or diseases episodes averted as a function of *p* and *q*, as estimated by a generalized additive model [62] with independent smooth terms for *p* and *q*. When one of *p* or *q* is varied, the other is held constant at 0.5. Gray bands indicate 95% confidence intervals.

Simulations conducted with the generic vaccine across the range of *p* also showed that proportions of cumulative infections and disease episodes averted were maximized (at 8.2% and 7.8% for VE_mean_=0.5, VE_serostatus_=0.075, VE_serotype_=0.075) when vaccine efficacy derived fully from protection against infection (*p*=1) (Fig. 10, left). Likewise, the lowest proportions of cumulative infections and disease episodes averted were observed when *p*=0 (0.25% and 2.9% for VE_mean_=0.5, VE_serostatus_=0.075, VE_serotype_=0.075). Because a much broader range of VE_mean_ was explored in these simulations than was represented by the range of *q* in simulations with the Dengvaxia^®^-like vaccine, the range of proportions of cumulative infections and disease episodes averted was much larger across the range of VE_mean_ than across the range of *q* (Fig. 10 vs. 9, right column). Differences in vaccine efficacy associated with serostatus and serotype had a more negligible effect on cumulative proportions of infections and disease episodes averted (Fig. 11).

**Figure 10.**
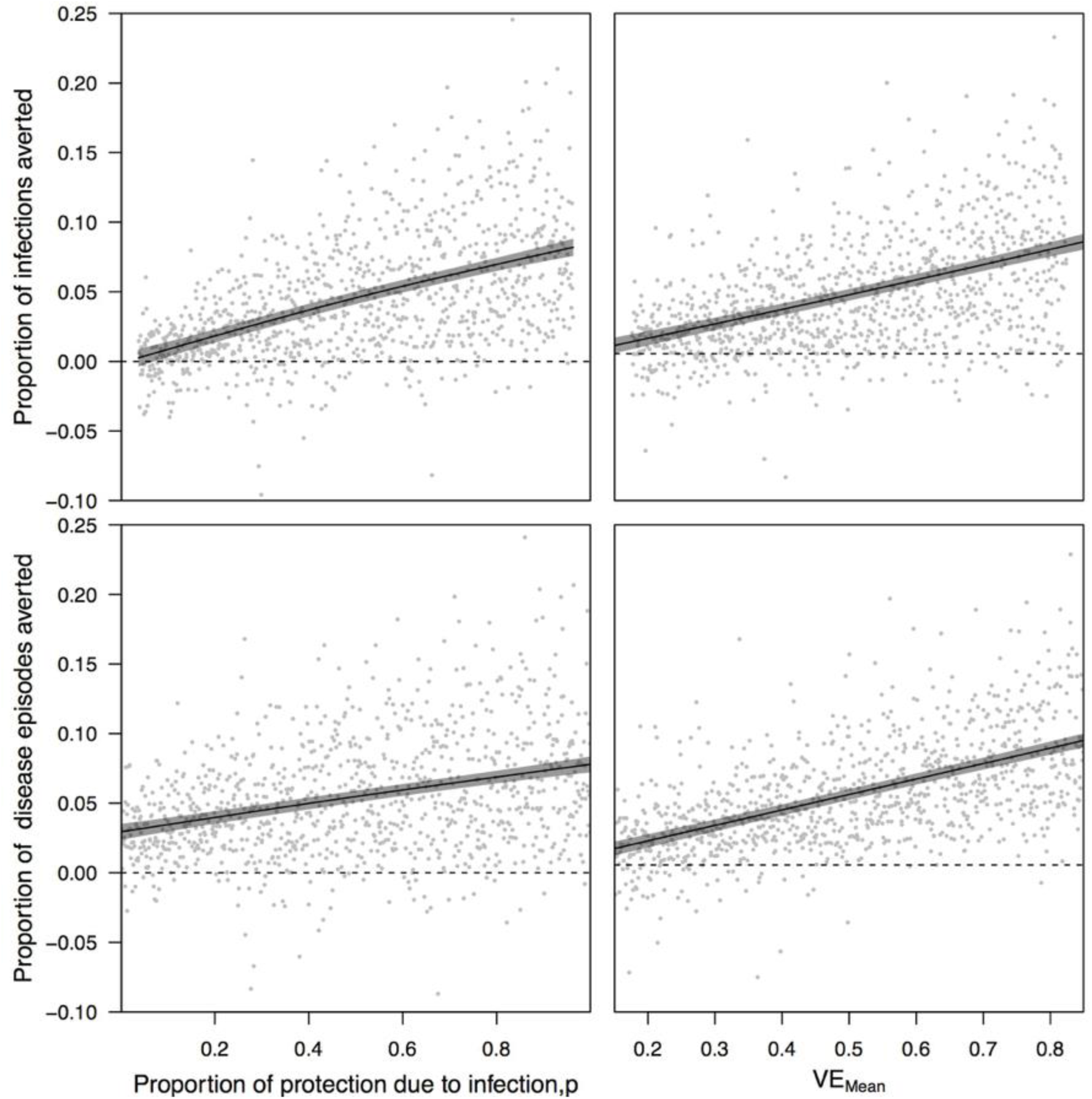
Impacts of vaccination assessed in 1,000 pairs of simulations with and without vaccination with the generic vaccine, with simulation pairs varying with respect to the proportion of vaccine efficacy due to protection from infection, *p*, (second column) and mean vaccine efficacy, VE_mean_, (right column). The proportion of cumulative infections (top row) and disease episodes (bottom row) averted were calculated as the number of each in the simulation without vaccination minus the number of each in the simulation with vaccination, both following 20 years of routine vaccination of 9-year olds at 80% coverage. Lines show the proportion of infections or diseases episodes averted as a function of each parameter varied on the x-axis, as estimated by a generalized additive model [62] with independent smooth terms for each parameter. When one parameter is varied, the other is held constant at the midpoint of its range, as are VE_serostatus_ and VE_serotype_. Gray bands indicate 95% confidence intervals.

**Figure 11.**
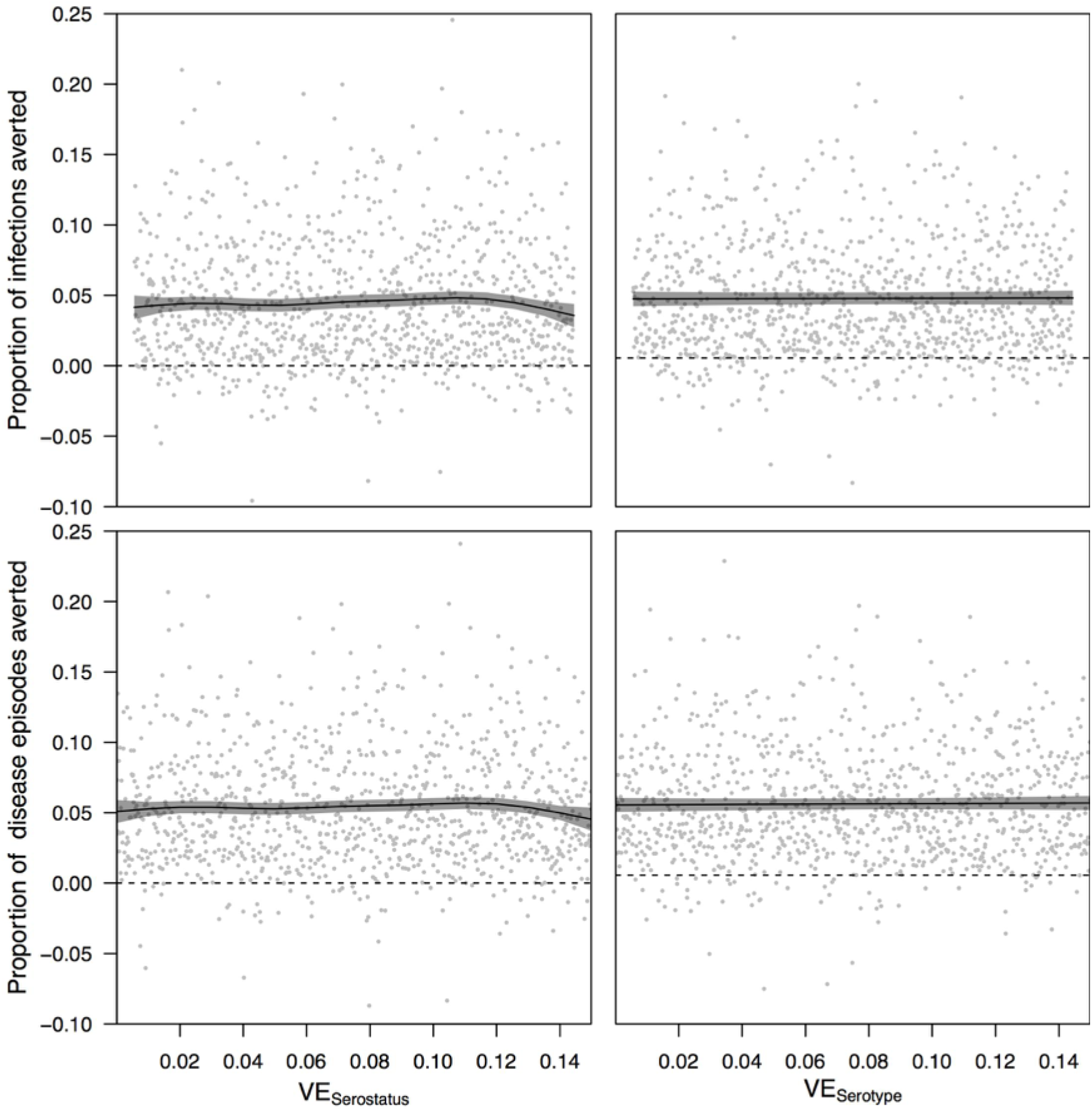
Impacts of vaccination assessed in 1,000 pairs of simulations with and without vaccination with the generic vaccine, with simulation pairs varying with respect to variation in vaccine efficacy associated with serostatus, VE_serostatus_, (second column) and variation in vaccine efficacy associated with serotype, VE_serotype_, (right column). The proportion of cumulative infections (top row) and disease episodes (bottom row) averted were calculated as the number of each in the simulation without vaccination minus the number of each in the simulation with vaccination, both following 20 years of routine vaccination of 9-year olds at 80% coverage. Lines show the proportion of infections or diseases episodes averted as a function of each parameter varied on the x-axis, as estimated by a generalized additive model [62] with independent smooth terms for each parameter. When one parameter is varied, the other is held constant at the midpoint of its range, as are *p* and VE_mean_. Gray bands indicate 95% confidence intervals.

In some simulations (Figs. 9-11), a negative proportion of infections or disease episodes was averted by vaccination (i.e., there were more cumulative infections or disease episodes in the simulation with vaccination). This was a result of chance differences in paired simulations with and without vaccination, due to the fact that random number seeds could maintain identical behavior with respect to some processes but not all.

## DISCUSSION

We developed an agent-based model for DENV transmission and applied it to questions regarding the impacts of uncertainty about different properties of dengue vaccines on projections of their public health impact. Our analysis was not intended to represent a comprehensive assessment of the suitability of Dengvaxia^®^ as a public health tool or to make a recommendation about its use. Instead, the value of this analysis is that it provides a theoretical assessment of the extent to which different sources of uncertainty about dengue vaccines might affect more detailed projections of public health impact. Our results indicate that future projections of dengue vaccine impact, applied either in specific settings or in generalities, should simulate across a range of multiple uncertainties about a vaccine’s properties, particularly regarding the degree to which efficacy derives from protection against infection versus protection against disease. Limiting such projections to only one set of assumptions along this continuum could result in the communication of recommendations to decision makers that convey a false sense of confidence in the presence of unrecognized uncertainty.

Our results were unambiguous in their suggestion that the projected public health impact of routine vaccination with Dengvaxia^®^ in nine-year olds is more sensitive to biological uncertainty about the degree of protection against infection than it is to statistical uncertainty about the numerical value of its efficacy against disease. This does not mean that the numerical value of vaccine efficacy is unimportant, but instead that uncertainty about its value following phase-III trials is low relative to uncertainty about the degree to which its efficacy derives from protection against infection or protection against disease. Echoing this, results from simulations using a generic dengue vaccine with mean efficacy ranging 0.15-0.85 show that the numerical value of vaccine efficacy is indeed important. Results from vaccine efficacy trials substantially narrow that range, because that is precisely what those trials are designed to do. At present, however, they are not designed to narrow uncertainty about the extent of infection blocking.

Based on our results and other theoretical studies [9,45], it is clear that knowledge of both efficacy against disease and efficacy against infection would be extremely valuable. Public health impact projections of tuberculosis vaccines came to similar conclusions that vaccines that protect against infection are more likely to have a long-term impact on reducing infection than are vaccines that emphasize protection from disease [45]. Even so, there are serious diagnostic limitations that impede the collection of data that would be required to estimate vaccine efficacy against infection, including extensive serological cross-reactivity among DENV and related viruses [46,47]. The same may also be true among naturally acquired and vaccine-induced immune responses. Similar challenges could plague the interpretation of Zika vaccine trial data, given that many Zika virus infections are mild or asymptomatic [48], generate immune responses that cross-react with other flaviviruses such as DENV [49], and may result in more severe disease among people with dengue antibodies [29,50].

The extent of differences that we observed due to biological uncertainty about the extent of protection against infection are a direct reflection of indirect effects of vaccination. Such effects have been predicted by models [51] and observed empirically [52] for a variety of diseases. One factor that is expected to affect the extent of indirect effects is coverage [53], with high coverage potentially compensating somewhat for low efficacy. For Dengvaxia^®^ or another dengue vaccine, it will be difficult to generalize about what coverage level might be appropriate or feasible across the range of settings where it could be applied. Another property that is expected to affect the extent of indirect effects are human-mosquito contact patterns [54]. In our model, we quantified contact patterns in a more realistic way than other DENV transmission models by leveraging published, data-driven models pertaining to human movement [14] and human-mosquito contact [20] in Iquitos, Peru. Realistically modeling contact patterns, vaccine coverage, and other parameters of relevance to indirect effects are all important considerations for public health impact projections, given the influence of indirect effects on cost-effectiveness calculations [55].

Although our model of vaccine efficacy is consistent with some findings from clinical trials of Dengvaxia^®^ [6], such as serostatus- and age-dependent efficacy against disease, there are other findings for which we did not account. One notable feature of the vaccine is the appearance of possible serotype-specific efficacy, with low efficacy against disease resulting from DENV-2 infection being of greatest concern [56]. A previous modeling analysis suggested that Dengvaxia^®^ may have a net positive impact on public health despite this shortcoming [57], and our results based on a generic dengue vaccine that did allow for serotype-specific efficacy indicated that this factor had a negligible effect. Other notable features of Dengvaxia^®^ that we have not considered pertain to protection against severe disease. In particular, to the extent that vaccination serves as a “primary-like” infection in seronegative vaccine recipients [38], the incidence of severe disease could increase as the proportion of seronegative vaccine recipients increases and transmission is lowered by indirect effects of vaccination [58]. Clinical trial data indicate, however, that whatever protection against severe disease the vaccine does afford may wane within a few years of vaccination [6]. It will be important for modeling analyses that seek to inform policy recommendations to account for the full complexity of the profile of the vaccine in question [59], but for the present analysis we chose to limit our assumptions about vaccine profile to those that are most pertinent to our driving questions and that can be readily informed with published data.

Although we expect that our qualitative results about the sensitivity of dengue vaccine impact projections to two forms of uncertainty about vaccine characteristics are applicable beyond Iquitos, we note that our quantitative projections of cumulative proportions of infections and disease episodes averted are not interpretable outside this setting. These numerical results could vary as a function of vaccination coverage, overall transmission intensity, and many other factors, similar to how estimates of vaccine efficacy can be context-dependent [60,61]. That said, our numerical projections based on a Dengvaxia^®^-like vaccine are in rough agreement with a study of vaccine impact based on an earlier version of our model and seven others (cumulative proportion of disease episodes averted ranged 6-25%, compared to our result of 14% assuming protection against infection) [43]. We hope that our model’s detailed representation of a well-studied, dengue-endemic population provides a tool for future studies to explore additional questions about vaccine impact in ways that acknowledge realistic variability in transmission patterns of the four distinct DENV serotypes.

## ACKNOWLEDGEMENT

This research made extensive use of computing resources provided by the Center for Research Computing at the University of Notre Dame.

## FUNDING

This research was supported by a grant from the US National Institutes of Health-National Institute of Allergy and Infectious Diseases (niaid.nih.gov) award 1P01AI098670-01A1 (to TWS) and by the Research and Policy for Infectious Disease Dynamics (RAPIDD) program of the Science and Technology Directory, Department of Homeland Security, and Fogarty International Center, National Institutes of Health (fic.nih.gov). DLS was supported by a grant from the National Institutes of Health (ICMER U19 AI089674), and VAPS was supported by a grant from the NIH Fogarty International Center (K01TW008414-01A1). TAP, RCR, and DLS were also supported by a grant from the Bill and Melinda Gates Foundation (OPP1110495) (gatesfoundation.org), and TAP and QAT received support from the Eck Institute for Global Health (globalhealth.nd.edu). The funders had no role in study design, data collection and analysis, decision to publish, or preparation of the manuscript.

## COMPETING INTERESTS

TAP, QAT, and GE receive partial support from a research contract from GlaxoSmithKline, which has a dengue vaccine candidate under development. GSK had no role in this study or in the preparation of this manuscript.

## SUPPORTING INFORMATION CAPTIONS

**Supporting Information 1. Detailed model description**.

**Supporting Information 2. Impact of number of exposures on the interpretation of vaccine efficacy**.

**Figure S1**. **Monthly, serotype-specific incidence of infection per capita, as estimated by Reiner et al. [16] (gray bands) and as reproduced by our calibrated model (colored bands)**. Bands show the range of values in which 95% of simulated values lie for a given serotype in a given month. These values were obtained under the assumption that the period of temporary cross-immunity is exponentially distributed with a mean of 180 days.

**Figure S2**. **Monthly, serotype-specific incidence of infection per capita, as estimated by Reiner et al. [16] (gray bands) and as reproduced by our calibrated model (colored bands)**. Bands show the range of values in which 95% of simulated values lie for a given serotype in a given month. These values were obtained under the assumption that the period of temporary cross-immunity is exponentially distributed with a mean of 360 days.

**Figure S3**. **Monthly, serotype-specific incidence of infection per capita, as estimated by Reiner et al. [16] (gray bands) and as reproduced by our calibrated model (colored bands)**. Bands show the range of values in which 95% of simulated values lie for a given serotype in a given month. These values were obtained under the assumption that the period of temporary cross-immunity is fixed at 180 days.

**Figure S4**. **Monthly, serotype-specific incidence of infection per capita, as estimated by Reiner et al. [16] (gray bands) and as reproduced by our calibrated model (colored bands)**. Bands show the range of values in which 95% of simulated values lie for a given serotype in a given month. These values were obtained under the assumption that the period of temporary cross-immunity is fixed at 360 days.

**Figure S5**. **Monthly, serotype-specific incidence of infection per capita, as estimated by Reiner et al. [16] (gray bands) and as reproduced by our calibrated model (colored bands)**. Bands show the range of values in which 95% of simulated values lie for a given serotype in a given month. These values were obtained under the assumption that the period of temporary cross-immunity is fixed at 686 days.

**Figure S6. Relationship between vaccine efficacy against disease (VE) and per-exposure protection, θ, for different values of the infection attack rate, over the period of a vaccine trial**.

## REFERENCES

1. Gubler DJ. The economic burden of dengue. Am J Trop Med Hyg. 2012;86: 743–744.

2. Achee NL, Gould F, Perkins TA, Reiner RC Jr, Morrison AC, Ritchie SA, et al. A critical assessment of vector control for dengue prevention. PLoS Negl Trop Dis. 2015;9: e0003655.

3. Morrison AC, Zielinski-Gutierrez E, Scott TW, Rosenberg R. Defining challenges and proposing solutions for control of the virus vector Aedes aegypti. PLoS Med. 2008;5: e68.

4. Vannice KS, Roehrig JT, Hombach J. Next generation dengue vaccines: A review of the preclinical development pipeline. Vaccine. 2015;33: 7091–7099.

5. WHO SAGE meeting of April 2016. World Health Organization; 2016; Available: http://www.who.int/immunization/sage/meetings/2016/april/en/

6. Hadinegoro SR, Arredondo-García JL, Capeding MR, Deseda C, Chotpitayasunondh T, Dietze R, et al. Efficacy and Long-Term Safety of a Dengue Vaccine in Regions of Endemic Disease. N Engl J Med. 2015;373: 1195–1206.

7. Grange L, Simon-Loriere E, Sakuntabhai A, Gresh L, Paul R, Harris E. Epidemiological risk factors associated with high global frequency of inapparent dengue virus infections. Front Immunol. 2014;5: 280.

8. Duong V, Lambrechts L, Paul RE, Ly S, Lay RS, Long KC, et al. Asymptomatic humans transmit dengue virus to mosquitoes. Proc Natl Acad Sci U S A. 2015;112: 14688–14693.

9. Rodriguez-Barraquer I, Mier-y-Teran-Romero L, Burke DS, Cummings DAT. Challenges in the Interpretation of Dengue Vaccine Trial Results. PLoS Negl Trop Dis 2013;7: e2126.

10. Morrison AC, Minnick SL, Rocha C, Forshey BM, Stoddard ST, Getis A, et al. Epidemiology of dengue virus in Iquitos, Peru 1999 to 2005: interepidemic and epidemic patterns of transmission. PLoS Negl Trop Dis. 2010;4: e670.

11. Stoddard ST, Wearing HJ, Reiner RC Jr, Morrison AC, Astete H, Vilcarromero S, et al. Long-term and seasonal dynamics of dengue in Iquitos, Peru. PLoS Negl Trop Dis. 2014;8: e3003.

12. Chao DL, Halstead SB, Elizabeth Halloran M, Longini IM Jr. Controlling Dengue with Vaccines in Thailand. PLoS Negl Trop Dis 2012;6: e1876.

13. Karl S, Halder N, Kelso JK, Ritchie SA, Milne GJ. A spatial simulation model for dengue virus infection in urban areas. BMC Infect Dis. 2014;14: 447.

14. Perkins TA, Garcia AJ, Paz-Soldán VA, Stoddard ST, Reiner RC Jr, Vazquez-Prokopec G, et al. Theory and data for simulating fine-scale human movement in an urban environment. J R Soc Interface. 2014;11.

15. United Nations, Department of Economic and Social Affairs, Population Division. World Population Prospects: The 2015 Revision - Special Aggregates, DVD Edition. 2015.

16. Reiner RC Jr, Stoddard ST, Forshey BM, King AA, Ellis AM, Lloyd AL, et al. Time-varying, serotype-specific force of infection of dengue virus. Proc Natl Acad Sci U S A. 2014;111: E2694–702.

17. Getis A, Morrison AC, Gray K, Scott TW. Characteristics of the spatial pattern of the dengue vector, Aedes aegypti, in Iquitos, Peru. Am J Trop Med Hyg. 2003;69: 494–505.

18. Grimm V, Berger U, Bastiansen F, Eliassen S, Ginot V, Giske J, et al. A standard protocol for describing individual-based and agent-based models. Ecol Modell. 2006;198: 115–126.

19. Grimm V, Berger U, DeAngelis DL, Polhill JG, Giske J, Railsback SF. The ODD protocol: A review and first update. Ecol Modell. 2010;221: 2760–2768.

20. Liebman KA, Stoddard ST, Reiner RC Jr, Perkins TA, Astete H, Sihuincha M, et al. Determinants of heterogeneous blood feeding patterns by Aedes aegypti in Iquitos, Peru. PLoS Negl Trop Dis. 2014;8: e2702.

21. Prothero RM. Disease and mobility: a neglected factor in epidemiology. Int J Epidemiol. 1977;6: 259–267.

22. Brady OJ, Johansson MA, Guerra CA, Bhatt S, Golding N, Pigott DM, et al. Modelling adult Aedes aegypti and Aedes albopictus survival at different temperatures in laboratory and field settings. Parasit Vectors. 2013;6: 351.

23. Reiner R, Stoddard S, Vazquez-Prokopec G, Astete H, Alex Perkins T, Sihuincha M, et al. Estimating the impact of city-wide Aedes aegypti population control: An observational study in Iquitos, Peru [Internet]. bioRxiv. 2018. p. 265751. doi:10.1101/265751

24. Harrington LC, Scott TW, Lerdthusnee K, Coleman RC, Costero A, Clark GG, et al. Dispersal of the dengue vector Aedes aegypti within and between rural communities. Am J Trop Med Hyg. 2005;72: 209–220.

25. Focks DA, Haile DG, Daniels E, Mount GA. Dynamic life table model for Aedes aegypti (Diptera: Culicidae): analysis of the literature and model development. J Med Entomol. 1993;30: 1003–1017.

26. Otero M, Solari HG, Schweigmann N. A stochastic population dynamics model for Aedes aegypti: formulation and application to a city with temperate climate. Bull Math Biol. 2006;68: 1945–1974.

27. Nguyen NM, Thi Hue Kien D, Tuan TV, Quyen NTH, Tran CNB, Vo Thi L, et al. Host and viral features of human dengue cases shape the population of infected and infectious Aedes aegypti mosquitoes. Proceedings of the National Academy of Sciences. 2013;110: 9072–9077.

28. Chan M, Johansson MA. The incubation periods of dengue viruses. PLoS One. 2012;7: e50972.

29. Clapham HE, Cummings DAT, Johansson MA. Immune status alters the probability of apparent illness due to dengue virus infection: Evidence from a pooled analysis across multiple cohort and cluster studies. PLoS Negl Trop Dis. 2017;11: e0005926.

30. Ellis AM, Garcia AJ, Focks DA, Morrison AC, Scott TW. Parameterization and sensitivity analysis of a complex simulation model for mosquito population dynamics, dengue transmission, and their control. Am J Trop Med Hyg. ASTMH; 2011;85: 257–264.

31. Nishiura H, Halstead SB. Natural History of Dengue Virus (DENV)—1 and DENV—4 Infections: Reanalysis of Classic Studies. J Infect Dis 2007; 195: 1007-1013.

32. Reich NG, Shrestha S, King AA, Rohani P, Lessler J, Kalayanarooj S, et al. Interactions between serotypes of dengue highlight epidemiological impact of cross-immunity. J R Soc Interface. 2013;10: 20130414.

33. Guagliardo SA, Morrison AC, Barboza JL, Requena E, Astete H, Vazquez-Prokopec G, et al. River boats contribute to the regional spread of the dengue vector Aedes aegypti in the Peruvian Amazon. PLoS Negl Trop Dis. 2015;9: e0003648.

34. Morrison AC, Gray K, Getis A, Astete H, Sihuincha M, Focks D, et al. Temporal and geographic patterns of Aedes aegypti (Diptera: Culicidae) production in Iquitos, Peru. J Med Entomol. 2004;41: 1123–1142.

35. Morrison AC, Sihuincha M, Stancil JD, Zamora E, Astete H, Olson JG, et al. Aedes aegypti (Diptera: Culicidae) production from non-residential sites in the Amazonian city of Iquitos, Peru. Ann Trop Med Parasitol. 2006;100 Suppl 1: S73–S86.

36. Reiner RC, Vazquez-Prokopec GM, Astete H, Perkins TA, Sihuincha M, Stancil JD, et al. Inferring the effect of vector control on Aedes aegypti in the face of spatio-temporal heterogeneity: an observational study in Iquitos, Peru.

37. Gordon NJ, Salmond DJ, Smith AFM. Novel approach to nonlinear/non-Gaussian Bayesian state estimation. IEE Proceedings F (Radar and Signal Processing). IET Digital Library; 1993;140: 107–113.

38. Guy B, Jackson N. Dengue vaccine: hypotheses to understand CYD-TDV-induced protection. Nat Rev Microbiol. 2016;14: 45–54.

39. Simmons CP. A Candidate Dengue Vaccine Walks a Tightrope. N Engl J Med. 2015;373: 1263–1264.

40. Team RC. R: A language and environment for statistical computing. R Foundation for Statistical Computing, Vienna, Austria. 2013. ISBN 3-900051-07-0; 2014.

41. Ewell M. Comparing methods for calculating confidence intervals for vaccine efficacy. Stat Med. 1996;15: 2379–2392.

42. Statistical Inference for Partially Observed Markov Processes [R package pomp version 1.4.1.1]. Comprehensive R Archive Network (CRAN); Available: https://cran.r-project.org/web/packages/pomp/index.html

43. Flasche S, Jit M, Rodríguez-Barraquer I, Coudeville L, Recker M, Koelle K, et al. The Long-Term Safety, Public Health Impact, and Cost-Effectiveness of Routine Vaccination with a Recombinant, Live-Attenuated Dengue Vaccine (Dengvaxia): A Model Comparison Study. PLoS Med. 2016;13: e1002181.

44. Halloran ME, Longini IM, Struchiner CJ, Longini IM. Design and analysis of vaccine studies. Springer; 2010.

45. Ziv E, Daley CL, Blower S. Potential public health impact of new tuberculosis vaccines. Emerg Infect Dis. 2004;10: 1529–1535.

46. Innis BL, Nisalak A, Nimmannitya S, Kusalerdchariya S, Chongswasdi V, Suntayakorn S, et al. An enzyme-linked immunosorbent assay to characterize dengue infections where dengue and Japanese encephalitis co-circulate. Am J Trop Med Hyg. 1989;40: 418–427.

47. Tang KF, Ooi EE. Diagnosis of dengue: an update. Expert Rev Anti Infect Ther. 2012;10: 895–907.

48. Duffy MR, Chen T-H, Hancock WT, Powers AM, Kool JL, Lanciotti RS, et al. Zika virus outbreak on Yap Island, Federated States of Micronesia. N Engl J Med. 2009;360: 2536– 2543.

49. Stettler K, Beltramello M, Espinosa DA, Graham V, Cassotta A, Bianchi S, et al. Specificity, cross-reactivity, and function of antibodies elicited by Zika virus infection. Science. 2016;353: 823–826.

50. Castanha PMS, Nascimento EJM, Braga C, Cordeiro MT, de Carvalho OV, de Mendonça LR, et al. Dengue Virus-Specific Antibodies Enhance Brazilian Zika Virus Infection. J Infect Dis. 2017;215: 781–785.

51. Van Effelterre T, Soriano-Gabarró M, Debrus S, Claire Newbern E, Gray J. A mathematical model of the indirect effects of rotavirus vaccination. Epidemiol Infect. 2010;138: 884–897.

52. Jordan R, Connock M, Albon E, Fry-Smith A, Olowokure B, Hawker J, et al. Universal vaccination of children against influenza: are there indirect benefits to the community? A systematic review of the evidence. Vaccine. 2006;24: 1047–1062.

53. Garnett GP. Role of herd immunity in determining the effect of vaccines against sexually transmitted disease. J Infect Dis. 2005;191 Suppl 1: S97–106.

54. Ma J, van den Driessche P, Willeboordse FH. The importance of contact network topology for the success of vaccination strategies. J Theor Biol. 2013;325: 12–21.

55. Brisson M, Edmunds WJ. Impact of model, methodological, and parameter uncertainty in the economic analysis of vaccination programs. Med Decis Making. 2006;26: 434–446.

56. Sabchareon A, Wallace D, Sirivichayakul C, Limkittikul K, Chanthavanich P, Suvannadabba S, et al. Protective efficacy of the recombinant, live-attenuated, CYD tetravalent dengue vaccine in Thai schoolchildren: a randomised, controlled phase 2b trial. Lancet. 2012;380: 1559–1567.

57. Rodriguez-Barraquer I, Mier-y-Teran-Romero L, Schwartz IB, Burke DS, Cummings DAT. Potential opportunities and perils of imperfect dengue vaccines. Vaccine. 2014;32: 514– 520.

58. Rodriguez-Barraquer I, Mier-y-Teran-Romero L, Ferguson N, Burke DS, Cummings DAT. Differential efficacy of dengue vaccine by immune status. Lancet. 2015;385: 1726.

59. Saadatian-Elahi M, Horstick O, Breiman RF, Gessner BD, Gubler DJ, Louis J, et al. Beyond efficacy: The full public health impact of vaccines. Vaccine. 2016;34: 1139–1147.

60. Lopman BA, Pitzer VE, Sarkar R, Gladstone B, Patel M, Glasser J, et al. Understanding reduced rotavirus vaccine efficacy in low socio-economic settings. PLoS One. 2012;7: e41720.

61. Gomes MGM, Gordon SB, Lalloo DG. Clinical trials: The mathematics of falling vaccine efficacy with rising disease incidence. Vaccine. 2016;34: 3007–3009.

62. Wood S. Generalized additive models: an introduction with R. CRC press; 2006.

63. Gillespie DT. Exact stochastic simulation of coupled chemical reactions. J Phys Chem. 1977;81: 2340–2361.

64. Glass K, Xia Y, Grenfell BT. Interpreting time-series analyses for continuous-time biological models—measles as a case study. J Theor Biol. 2003;223: 19–25.

65. Sabin AB. Research on dengue during World War II. Am J Trop Med Hyg. 1952;1: 30–50.

66. UNdata [Internet]. [cited 30 Apr 2016]. Available: http://data.un.org

67. Perkins TA, Siraj AS, Ruktanonchai CW, Kraemer MUG, Tatem AJ. Model-based projections of Zika virus infections in childbearing women in the Americas. Nat Microbiol. 2016; 1: 16126.

68. Olkowski S, Forshey BM, Morrison AC, Rocha C, Vilcarromero S, Halsey ES, et al. Reduced risk of disease during post-secondary dengue virus infections. J Infect Dis. 2013; 208: 1026–1033.

69. Forshey BM, Reiner RC, Olkowski S, Morrison AC, Espinoza A, Long KC, et al. Incomplete Protection against Dengue Virus Type 2 Re-infection in Peru. PLoS Negl Trop Dis. 2016;10: e0004398.

